# WFA-GPU: Gap-affine pairwise alignment using GPUs

**DOI:** 10.1101/2022.04.18.488374

**Authors:** Quim Aguado-Puig, Max Doblas, Christos Matzoros, Antonio Espinosa, Juan Carlos Moure, Santiago Marco-Sola, Miquel Moreto

## Abstract

**Motivation:** Advances in genomics and sequencing technologies demand faster and more scalable analysis methods that can process longer sequences with higher accuracy. However, classical pairwise alignment methods, based on dynamic programming (DP), impose impractical computational requirements to align long and noisy sequences like those produced by PacBio, and Nanopore technologies. The recently proposed WFA algorithm paves the way for more efficient alignment tools, improving time and memory complexity over previous methods. However, high-performance computing (HPC) platforms require efficient parallel algorithms and tools to exploit the computing resources available on modern accelerator-based architectures.

**Results:** This paper presents the WFA-GPU, a GPU (Graphics Processing Unit)-accelerated tool to compute exact gap-affine alignments based on the WFA algorithm. We present the algorithmic adaptations and performance optimizations that allow exploiting the massively parallel capabilities of modern GPU devices to accelerate the alignment computations. In particular, we propose a CPU-GPU co-design capable of performing inter-sequence and intra-sequence parallel sequence alignment, combining a succinct WFA-data representation with an efficient GPU implementation. As a result, we demonstrate that our implementation outperforms the original multi-threaded WFA implementation between 1.5-7.7× and up to 17× when using heuristic methods on long and noisy sequences. Compared to other state-of-the-art tools and libraries, the WFA-GPU is up to 29× faster than other GPU implementations and up to four orders of magnitude faster than other CPU implementations.

**Availability:** WFA-GPU code and documentation are publicly available at https://github.com/quim0/WFA-GPU.

## 1 Introduction

Pairwise sequence alignment is a fundamental building block in many tools and libraries used in genomics and bioinformatics. In particular, it plays a critical role for methods like read mapping (Li, 2013; Marco-Sola *et al*., 2012), de-novo genome assembly (Simpson *et al*., 2009; Koren *et al*., 2017), variant calling (McKenna *et al*., 2010; Rodríguez-Martín *et al*., 2017), and many others (Durbin *et al*., 1998; Jones *et al*., 2004).

Consequently, sequence alignment algorithms have been extensively studied over the last 40 years, introducing multiple strategies like dynamic programming (DP) (Sellers, 1980; Ukkonen, 1985), automata (Baeza-Yates, 1992; Wu and Manber, 1992; Navarro, 1997), and bit-parallelism techniques (Myers, 1999; Baeza-Yates, 1989). Nonetheless, these algorithms are bound by the quadratic time and memory requirements on the sequence length. Thus, the use of classical alignment algorithms becomes impractical as the input sequences increase in length. Many variations and optimizations have been proposed over the years to overcome these limitations. These solutions include techniques such as banded approaches that only compute a portion of the DP-matrix (Suzuki and Kasahara, 2017), data-layout organizations that allow using SIMD instructions (Rognes and Seeberg, 2000; Farrar, 2007; Wozniak, 1997), bit-packed encodings (Myers, 1986, 1999), and other methods (Ukkonen, 1985; Zhao *et al*., 2013). Nevertheless, all these proposals retain the quadratic requirements and fail to scale for long sequences.

Recently, the Wavefront Alignment (WFA) algorithm (Marco-Sola *et al*., 2021) was proposed. The WFA is able to compute the exact alignment between two sequences using gap-affine penalties. In essence, the WFA algorithm computes partial alignments of increasing score *s* until the optimal alignment is found. The WFA algorithm takes advantage of homologous regions between sequences to accelerate the alignment process. As a result, the WFA algorithm largely outperforms other state-of-the-art methods, requiring *O*(*ns*) time and *O*(*s*^2^) memory (where *n* is the length of the sequence and *s* is the optimal alignment score).

Notwithstanding, the ever-increasing yields and read lengths produced by sequencing machines challenge the scalability of current sequence alignment methods. In particular, modern sequencing technologies, like PacBio or Oxford Nanopore, can produce sequences more than 100× longer than those produced by classical Illumina sequencers at a fraction of the cost. The adoption of these technologies calls for the development of faster and more scalable alignment solutions (Petersen *et al*., 2019). So imperative is the need to scale to larger volumes of genomic data, that the adoption of HPC solutions has become more and more frequent. In particular, GPUs have been widely adopted as hardware accelerators in many scientific applications (Hwu, 2011; Owens *et al*., 2008; Chacón *et al*., 2014; Lin *et al*., 2017) as they provide higher computational throughput and memory bandwidth compared with traditional multi-core processors. In this context, alignment algorithms and tools need to address the efficient exploitation of these hardware accelerators to keep up with the pace of modern sequence data production.

This paper presents WFA-GPU, a GPU-accelerated implementation of the WFA algorithm for exact gap-affine pairwise sequence alignment. We describe the adaptations performed on the WFA algorithm to exploit the massively parallel capabilities of modern GPU architectures. In particular, our proposal combines inter-sequence and intra-sequence parallelism to speed up the alignment computation. Moreover, we propose a succinct backtrace encoding to reduce the overall memory consumption of the original WFA algorithm. Additionally, we present a heuristic variant of the WFA-GPU that further improves its performance, achieving nearly the same accuracy as the original exact WFA algorithm. We characterize the performance of our implementation and present the different performance trade-offs of our solution. As a result, we demonstrate that our implementation outperforms other GPU and CPU state-of-the-art libraries and tools for sequence alignment.

The rest of the paper is structured as follows. Section 2 presents the definitions and methods of our proposal. Section 3 shows the experimental results, comparing the performance of our method against other state-of-the-art implementations on both CPU and GPU systems. Finally, Section 4 presents a discussion on the methods presented and summarizes the contributions and impact of this work.

## 2 Methods

### 2.1 Wavefront Pairwise Alignment

Let the query *q* = *q*_0_*q*_1_…*q*_*n*−1_ and the text *t* = *t*_0_*t*_1_…*t*_*m*−1_ be strings of length *n* and *m*, respectively. Similarly, let *q*_*i*…*j*_ denote a substring of *q* from the *i*-th to the *j*-th character (both included). Also, let {*x, o, e*} be the set of gap-affine penalties, where *x* is the mismatch cost and the gap cost is expressed as the linear function *g*(*l*) = *o* + *l* · *e* (where *l* is the length of the gap).

In essence, the WFA algorithm computes partial alignments of increasing score until an alignment with score *s* reaches coordinate (*n, m*) (i.e., the end of the DP matrix). This way, the algorithm is able to compute the optimal score *s* and retrieve the optimal alignment by tracing back the alignment operations (i.e., {*M, X, I, D*} for match, mismatch, insertion, and deletion) that led to the solution.

Let 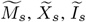, and 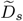 be the wavefront components that describe partial alignments of score *s* that end with a match, mismatch, insertion, and deletion, respectively. In general, we denote 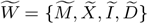 as the set of wavefront components. We define 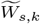 as the farthest reaching point of score *s* on diagonal *k*. That is, 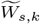 denotes the coordinate 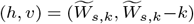 in the DP matrix that is farthest in the diagonal *k* with score *s*. Thus, a wavefront 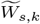 is a vector containing the farthest reaching points with score *s* on each diagonal *k*. In (Marco-Sola *et al*., 2021), the authors prove that the farthest reaching points of 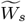 can be computed using the wavefronts with score *s* − *o* − *e, s* − *e*, and *s* − *x*, using Eq. 1 (where *LCP*(*v,w*) is the length of the longest common prefix between two string *v* and *w*). Note that the wavefront component 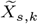 can be inferred using 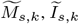, and 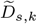, and we can avoid storing it.

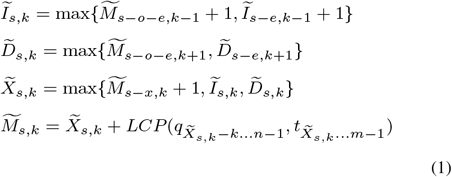

Starting with 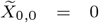, the WFA algorithm progressively computes wavefronts of increasing score. For a given score *s*, it first increases each wavefront offset 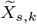 according to the number of matching characters along the diagonal *k* (i.e., computing 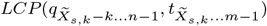 for every diagonal *k*). Then, the algorithm computes the next wavefronts with score *s* + 1 using Eq. 1 and the previously computed wavefronts. This process iterates until a wavefront 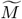 reaches the bottom-right cell (*n, m*) of the DP matrix (i.e., 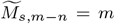). Hence, the optimal alignment score is *s* and the alignment operations can be retrieved by tracing back the wavefronts that originated the farthest reaching point 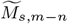.

In the worst case, the WFA algorithm requires computing *s* wavefronts of increasing length 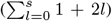 and compute the *LCP()* for each wavefront offset. Nevertheless, each diagonal offset cannot be incremented more than the length of the sequence. Thus, the WFA algorithm requires *O*(*ns* + *s*^2^) time (*O*(*ns*) if the sequences share similarities, as *s << n*) and *O*(*s*^2^) memory in the worst case.

### 2.2 Graphics Processing Units

GPUs have rapidly emerged as successful hardware accelerators in the HPC community for scientific applications. GPUs are massively parallel devices containing multiple throughput-oriented processing units called Stream Multiprocessors (SMs). Each SM can host over a thousand concurrent threads, and each clock cycle, using aggressive fine-grained multithreading techniques, can start executing hundreds of instructions on multiple SIMD cores. A CPU-GPU co-design involves determining a computationally intensive function or region of a program (i.e., computation kernel) to be offloaded to the GPU. To exploit GPU parallel computing capabilities, applications must launch tens of thousands of threads, grouped in thread blocks, across all available SMs. Threads within a block can cooperate (e.g., via synchronization primitives, registers, or shared memory) to solve a given computational kernel. To maximize performance, GPUs rely on exploiting high degrees of parallelism, much higher than those of regular CPU multi-cores.

Each SM has fast on-chip memories with a relatively small capacity. They are distributed among compiler-managed register memory (up to 256KB), processor-managed L1 cache, and programmer-managed shared memory (up to 100KB for the total sum of L1 and shared memory). The SMs get their data from an off-chip RAM memory of several GBs capable of delivering bandwidths of over 600 GB/s. An L2 cache holds several MBs of the RAM memory contents and offers 3-5 times higher bandwidth.

The peak computational throughput of GPUs is much higher than its peak memory bandwidth. Therefore, doing very few memory accesses per arithmetic operation is paramount to achieving good GPU utilization. The use of fast on-chip memories (registers, L1, and shared memory) can alleviate this problem. Still, there is a critical compromise between the number of concurrent threads hosted by each SM and the amount of data each thread can store in fast memories. Allocating too many threads per SM results in more threads competing for the relatively small fast memories, increasing the number of data accesses to slow memory, and performance may suffer. Reducing the number of threads per SM can improve memory performance, but at the price of reducing parallelism and the ability to hide long execution and memory latencies, potentially decreasing performance.

#### Algorithm 1: WFA-GPU parallel algorithm

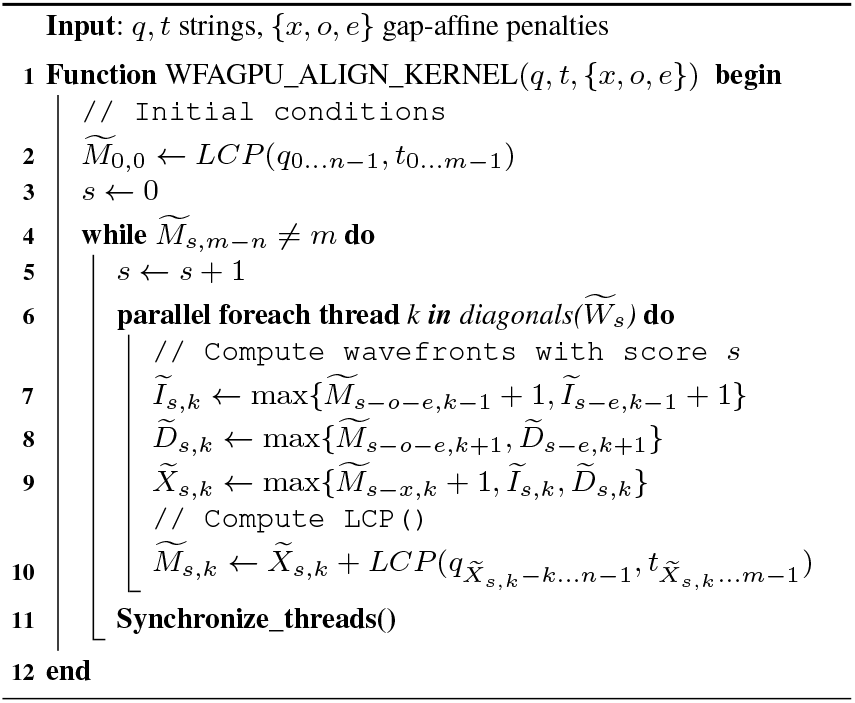

### 2.3 WFA-GPU

In this section, we present the parallel WFA-GPU algorithm, the main performance challenges, and effective solutions to mitigate these problems. This way, Section 2.3.1 presents an overview of the parallel algorithm and its mapping to the GPU computing architecture. Afterwards, we discuss the principal performance limitations of its implementation on GPU.

First, we show that the memory requirements grow quadratically as the alignment error increases, limiting the scalability of the implementation when aligning tens of thousands of sequences in parallel. To alleviate this problem, in Sections 2.3.2 and 2.3.3, we present an efficient GPU memory management strategy and an algorithmic technique to reduce the overall memory used by the WFA on GPU.

Second, we argue that the number of operations to compute successive wavefronts becomes a limiting factor when aligning large and noisy sequences. In Section 2.3.4, we present a strategy to accelerate the *LCP()* computation and, in Section 2.3.5, we propose a GPU-aware heuristic extension to reduce the volume of computations performed on the GPU.

Finally, in Section 2.3.6, we present a CPU-GPU co-design, which allows overlapping the execution of GPU kernels, CPU tasks, and data transfers (between the memories of the two devices) to improve the overall performance of the implementation.

#### 2.3.1 WFA-GPU parallel algorithm

Classical parallel solutions to the sequence alignment problem are based on exploiting inter-sequence parallelism (i.e., letting each GPU thread compute a different alignment). However, as parallelism increases, aligning a large number of sequences simultaneously requires unfeasible amounts of memory. Moreover, differences in the workload (e.g., sequence length and alignment error) cause variations in the execution flow of each alignment, generating thread divergence. Alternatively, it is feasible to exploit parallelism within a single alignment task (i.e., intra-sequence parallelism). However, a single alignment rarely allows exploiting the massive amounts of parallel resources available on modern GPUs. Our proposal is to combine both parallel strategies (inter- and intra-parallelism) to compute multiple alignments on concurrent thread blocks, each using several threads cooperatively to calculate a single alignment.

The WFA algorithm depicts a simple computational pattern to calculate each sequence alignment. Eq. 1 shows that a wavefront 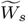 is computed using only wavefronts 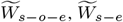, and 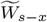. More importantly, each diagonal offset 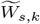 can be computed independently. Fig. 1 illustrates in detail the parallel computation of a given wavefront *s* using multiple GPU threads, exploiting intra-sequence parallelism 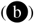. At the same time, other alignments can be computed by different thread blocks on the GPU using inter-sequence parallelism 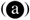.

**Fig. 1:**
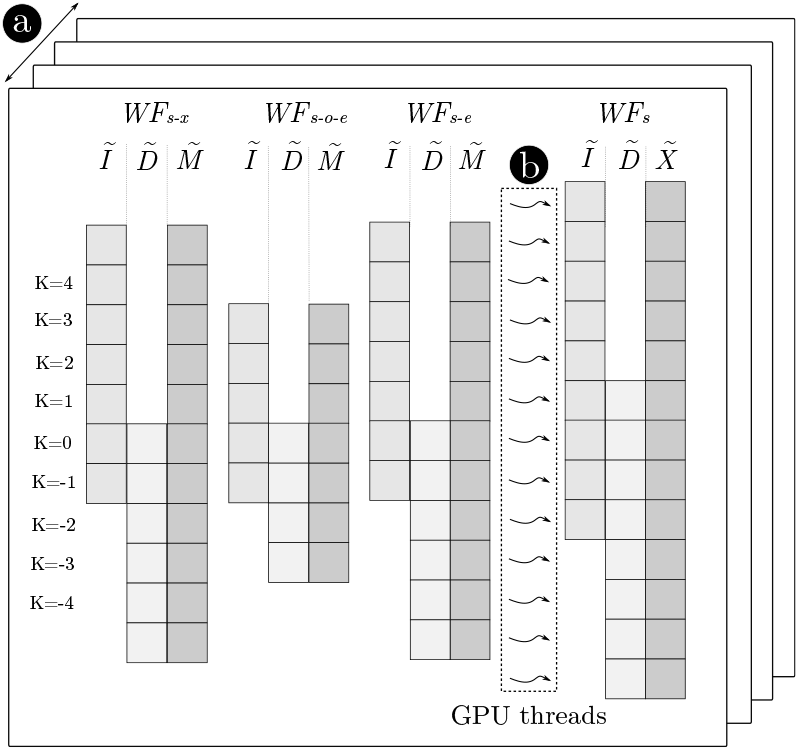
Multiple alignments are computed concurrently on the GPU, exploiting inter-sequence parallelism 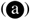. At the same time, multiple GPU threads compute different diagonals of a given wavefront (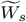) in parallel, exploiting intra-sequence parallelism 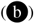.

Algorithm 1 presents the high-level pseudocode of the WFA-GPU. For each alignment, multiple threads cooperate to compute consecutive wavefronts. In particular, for every score *s* and diagonal *k*, each GPU thread in the block computes 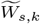 independently (lines 7-10). After every diagonal of wavefronts 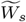 is computed, GPU threads synchronize (line 11) and proceed to compute the following wavefronts 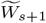. This process is repeated until the optimal alignment is found (line 4).

Each GPU thread concurrently undertakes the computation of the data item of a different diagonal offset, applying Eq. 1 and using the previously computed wavefronts and the *LCP()* function. Notably, as the algorithm progresses, wavefronts become increasingly larger and the potential parallelism of the problem grows. For large and noisy sequences, the problem becomes embarrassingly parallel, allowing to perform up to 2*s*+1 parallel computations for each 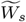. Such a highly parallel algorithm is suitable for modern GPUs.

#### 2.3.2 Alignment Scheduling and GPU Memory Management

A simple and naive implementation would spawn a thread block per each WFA alignment offloaded to the GPU. However, each WFA alignment kernel requires GPU memory to store all the intermediate wavefronts. It is not feasible to reserve GPU memory for every alignment in advance when processing tens of thousands of sequence alignments. However, it is possible to request an upper bound of the total WFA memory required for a number of alignments that can be processed in parallel in the GPU at the same time.

Thus, our implementation creates a pool of outstanding WFA alignments and allocates memory for as many alignment blocks as can be processed simultaneously on the GPU. Then, an alignment scheduler assigns WFA alignments to thread blocks. Whenever a thread block finishes an alignment, it requests another from the alignment pool until all alignments offloaded to the GPU have been completed.

Nevertheless, having to allocate GPU memory beforehand forces to estimate the maximum memory required by each WFA alignment in advance. For that, our method establishes a configurable upper bound on the required memory based on a conservative estimation of the maximum error rate (i.e., 10% of sequence length by default on our example tool). Nonetheless, some alignments may override initial estimations and require more memory. For those cases, the WFA-GPU implements a rescue mechanism that returns the alignment to the CPU to be computed using the original WFA algorithm. In practice, when aligning long and noisy sequences (like those produced by PacBio or Nanopore Technologies) the amount of rescued alignments is below 0.2%. Furthermore, the computation of the rescued alignments can be performed in the host CPU; meanwhile, the GPU is computing other alignments, as described in Section 2.3.6.

Although modern GPUs are equipped with large DRAM memories, accesses to global memory are relatively slow and can potentially reduce the performance of GPU applications. To take advantage of fast on-chip memories and minimize the latency of global memory accesses in the GPU, our implementation allocates the most frequently accessed wavefront diagonals (i.e. the central diagonals) of wavefronts in the shared memory. This way, the WFA-GPU benefits from fast on-chip memory access to the elements of the central diagonals.

#### 2.3.3 Piggybacked Backtrace Operations

For an alignment with optimal score *s*, the WFA algorithm requires storing all the intermediate wavefronts up to 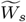 to be able to retrieve the alignment path (a.k.a. CIGAR) during the final backtrace step. However, alignments with a large nominal number of errors require a non-negligible amount of memory. That is, an upper-bound of 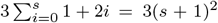 wavefronts offsets, consuming up to 12(*s* + 1)^2^ bytes per alignment. These memory requirements become impractical when aligning multiple noisy sequences in parallel, even for modern GPUs equipped with large amounts of global memory.

To reduce the memory consumption, our method piggybacks the backtrace operations (i.e., **X, I**, and **D**) to the wavefronts as they are being computed. Using only two bits, each backtrace operation is encoded in a bitmap stored for every diagonal of the wavefront. Therefore, for a given score *s* and diagonal *k*, our method stores a bitmap with the alignment operations required to reach 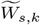. It follows that the bitmap associated with 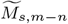 contains the optimal alignment’s backtrace. Note that these bitmaps do not grow regularly, as a different number of backtrace operations may have the same score (e.g., multiple insertions may be equivalent to a mismatch).

In practice, our implementation uses 32-bit bitmap words to store backtrace operations (i.e., BT-block). Once a BT-block of a diagonal is full and cannot encode more backtrace operations (❶ on Fig. 2), it is offloaded to a global backtrace buffer (❷) (i.e., BT-buffer). Each BT-block stores an index to the previous BT-block (❸) in the chain that allows retrieving the complete alignment backtrace associated with any 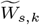 offset. A complete alignment backtrace is recovered by traversing the linked BT-blocks starting from the last one.

**Fig. 2:**
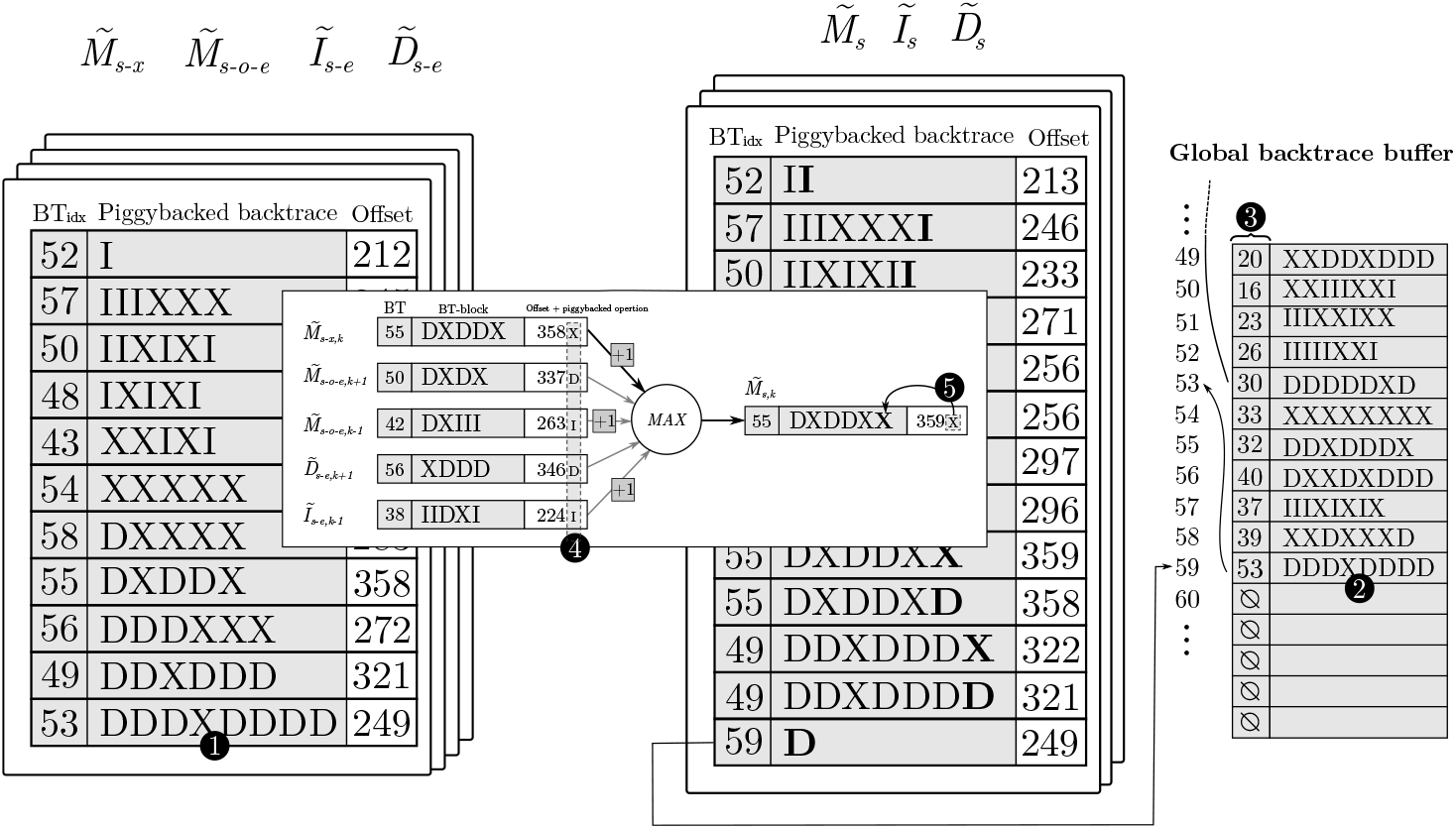
Illustration of the piggybacked backtrace strategy and data-layout organization. From left to right of the figure, we show the source wavefronts (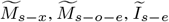, and 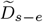) and how they are combined to generated wavefronts at score *s*. Each diagonal has a BT-block (shaded in grey) and an offset. At the center, we detail the process of computing Eq. 1 for a single diagonal and the piggyback of the corresponding backtrace operation. At the bottom, the global BT-buffer is depicted, where each slot represents a BT-block (displayed vertically for better readability of the figure).

The computation of each backtrace operation is coupled with the computation performed in Algorithm 1 to generate each diagonal offset. For that, the corresponding backtrace operation (i.e., **X, I**, and **D**) is piggybacked to each source wavefront offset (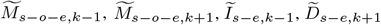, and 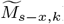) in Eq. 1, using the two less significant bits (❹). Then, as a byproduct of the computation of the maximum offset, the corresponding backtrace operation is found piggybacked, and is added to the current BT-block (❺). In practice, this strategy turns out to be computationally lightweight.

Note that BT-blocks only contain edit operations (i.e. **X, I, D**) and not the matches in between. To retrieve the complete alignment CIGAR, the algorithm needs to compute any missing matches between backtrace operations. Nonetheless, this is a remarkably simple operation. Using the same *LCP()* function presented earlier, the algorithm computes matches until a mismatch is found. Then, it adds the following backtrace operation and proceeds again to compute the *LCP()*. This process halts when all the backtrace operations from the chain of BT-blocks have been processed.

Overall, the piggyback strategy effectively reduces the memory consumed by the wavefronts to 4 bits per entry (accounting for the BT-block indices). Compared to storing the raw wavefront offsets as the original WFA does (i.e., 4 bytes per entry), this strategy represents an 8× reduction.

#### 2.3.4 Bit-Parallel Sequence comparison using Packed DNA Sequences

WFA’s execution time is dominated by the computation of the *LCP()* function (Eizenga and Paten, 2022). A naive implementation would compare sequences character by character until a non-matching character is found. This approach not only executes a non-negligible amount of instructions per *LCP()* call but also creates divergence across threads computing the same alignment. That is, each GPU thread within a block performs a different number of comparisons depending on the characters being compared. Because GPU threads execute in groups of 32 threads in lock-step mode, divergent execution (i.e., variable-iterations loops) forces idle threads to wait until all threads have finished iterating.

To alleviate this problem, we propose a bit-packed encoding of DNA sequences using 2 bits per base. The encoding turns out to be remarkably simple, as the ASCII representation of each base has two unique bits on position 1 and 2 (i.e., A=1000**00**1, C=1000**01**1, G=1000**11**1, T=1010**10**0). This process is also well suited for GPU execution, as each character can be computed in parallel. Sequence packing is done at runtime, just before the WFA-GPU alignment kernel starts. Using a bit-packed representation, our implementation compares blocks of 16 bases at once using 32-bit operations. This strategy reduces execution divergence and, most importantly, the total number of instructions executed, which translates into faster execution times.

#### 2.3.5 GPU-aware Approximated Wavefront Alignment

For some applications, the exact computation of the alignment may be unnecessary, and a reasonable approximation of the optimal solution may suffice (e.g., pre-filtering (Alser *et al*., 2019, 2017) and clustering (Zorita *et al*., 2015; Zou *et al*., 2020) applications). For that, heuristic strategies can reduce the number of computations required, improving performance at the expense of a potential loss of accuracy. In addition to the exact WFA-GPU algorithm, we propose a heuristic extension that improves the performance of the exact algorithm achieving nearly the same accuracy. Our heuristic strategy draws inspiration from the original WFA-Adapt (Marco-Sola *et al*., 2021) heuristic and other adaptive techniques (Suzuki and Kasahara, 2017). Unlike previous approaches, our strategy is tailored to the architecture and resources available in the GPU and exploits the GPU parallelism to minimize the overhead of executing the heuristic.

Our heuristic strategy employs a fixed-size wavefront of length *β*. Initially, wavefronts are centered around the main diagonal. Every *λ* steps, the algorithm computes the most promising diagonal. That is, the diagonal closest to the target cell (the bottom-right corner of the DP table). Afterwards, newly computed wavefronts are centered around the most promising diagonal, focusing the wavefront computation towards the most promising partial alignment. As a result, the heuristic avoids the computation of wavefront diagonals that are left behind (unlikely to lead to the optimal solution).

Selecting adequate values of *β* and *λ* can have an impact on performance and accuracy. Note that a small *β* can lead to significant accuracy loss. Similarly, a small *λ* could lead to overheads computing the heuristic. As opposed, large values of *β* and *λ* could render the heuristic ineffective. Our implementation selects a value of *β* such that all the wavefronts can fit into the fast on-chip shared memory of the GPU. Regarding *λ*, we observe that values *λ* ≤ 100 perform similarly in terms of accuracy. Moreover, to alleviate the computational burden when *λ* is small, our implementation uses a thread-cooperative strategy. For that, threads within a warp cooperate to find the most promising diagonal and center the wavefront around it.

#### 2.3.6 CPU-GPU Co-design System

The WFA-GPU implements a CPU-GPU co-design that allows the simultaneous execution of GPU computations overlapped with data transfers and CPU alignment rescue. To maximize performance, our implementation offloads batches containing multiple alignments to the GPU. For that, input sequences from a batch have to be transferred to the device. To minimize GPU idle times, our implementation makes asynchronous kernel launches, allowing overlapping data transfers with GPU computations. That is, while the GPU is computing the alignments for a given batch, the sequences of the following batch are being copied to the device. As a result, latencies due to transfer times are effectively hidden and overlap with useful GPU computations.

Furthermore, the asynchronous implementation of WFA-GPU allows employing idle CPU time to rescue alignments returned by the GPU. As explained in Section 2.3, a small percentage of alignments may not be aligned in the GPU due to exceeding memory requirements. For those few cases, the implementation overlaps the CPU WFA execution with GPU computations and data transfers.

## 3 Results

We evaluated the performance of the WFA-GPU, together with other state-of-the-art CPU and GPU tools for sequence alignment. In Section 3.1, we present the system specifications and datasets used. In Section 3.2, we present an evaluation using simulated datasets. Finally, in Section 3.3, we present the experimental results using real datasets.

### 3.1 Experimental Setup

For the experimental evaluation, we select simulated and real datasets. For the simulated datasets, we generate synthetic pairs of sequences of 150, 1000, and 10,000 bases aligning with an average edit-error of 2%, 5%, and 10% differences. For the evaluation using real datasets, we select publicly available datasets representative of current sequencing technologies (see Table S1). The target sequences are retrieved from mapping the source sequences against GRCh38 using Minimap2 (Li, 2018) and default parameters.

To compare the performance of the WFA-GPU, we select other sequence alignment libraries and tools representative of the state-of-the-art on both CPU and GPU devices. For the GPU tools comparison, we select the library GASAL2 (Ahmed *et al*., 2019), ADEPT (Awan *et al*., 2020) and two NVIDIA libraries (NVBio (Pantaleoni J, 2015) and CudaAligner (cud, 2022) from Clara Parabricks Genomeworks).

Unfortunately, we were unable to include the recently presented GPU aligners Logan (Zeni *et al*., 2020) and GenASM (Lindegger *et al*., 2022) due to inadequacy to perform basic pairwise alignment and the unavailability of the source code, respectively. As for the CPU tools, we selected the most widely-used and efficient libraries available to date. That is, Seqan (Döring *et al*., 2008), Parasail (Daily, 2016), Edlib (Šošić and Šikić, 2017), and KSW2 (Suzuki and Kasahara, 2018). Naturally, we also include the original WFA implementation (WFA2-lib (wfa, 2022)) in the comparison. Note that Edlib and CudaAligner can only compute the edit distance alignment (a much simpler problem compared to computing gap-affine alignments). Regardless, we include them in the comparison as an interesting point of reference. In an attempt to evaluate the recall of these tools using gap-affine scores, we re-scored the reported CIGAR using gap-affine penalties and compared it with the optimal score. ADEPT computes the local alignment of two sequences, as we compute global alignment, ADEPT can not be compared with WFA-GPU in terms of accuracy. This is indicated as *not-comparable* (n/c) in the tables.

All the experiments are executed using a 10-core Intel Xeon-W2155 (3.3GHz) processor equipped with 126GB of memory and a NVIDIA GeForce 3080 with 10GB of memory. Moreover, all CPU executions are performed in parallel using the 10 physical cores available in the platform. All GPU execution times include CPU-GPU data transfer, alignment, backtrace, and CIGAR generation time.

### 3.2 Evaluation on Simulated Data

Table S2 shows time (in seconds) and recall (percentage of sequences for which the optimal alignment was correctly reported) for the alignment executions using simulated datasets.

Considering the alignment of short sequences (i.e., ∼150bps), NVBio outperforms all other tools at the expense of a significant loss in alignment accuracy as the alignment error increases. WFA-GPU is between 1.9 and 3 times faster than the best CPU time obtained. GASAL2 shows good performance (as it is tailored to short sequence alignment). Because dynamic-programming approaches are error-independent, GASAL2 outperforms our implementation when the error is high. The other GPU aligners, ADEPT and CudaAligner, are one order of magnitude slower than WFA-GPU and GASAL2. For these short-sequence datasets, using the WFA-GPU approximated alignment does not provide any significant benefit because the error is very small.

For medium-length sequences (∼1Kbps), GPU implementations either fail due to execution errors (like NVBio and ADEPT) or obtain a recall below 10%. Only GASAL2 remains competitive when the error increases, but at a significantly low accuracy (52.4%). Compared to CPU implementations, WFA-GPU executes 3.2-5.8× faster than the original WFA and up to 5500× faster than other libraries. Using WFA-GPU approximated alignment, we obtain an additional speedup of up to 3× when computing the optimal alignment path (backtrace).

Experiments aligning long simulated sequences (i.e., ∼10Kbps) turn out to be the most challenging for most GPU tools. All other GPU implementations either fail (i.e., ADEPT and NVBio), give incorrect results (i.e., CudaAligner), or have significantly low recall (i.e. GASAL2 with less than 50% accuracy). WFA-GPU is the only GPU implementation that can scale to long sequences reporting the optimal alignment result. When compared to the original WFA, WFA-GPU executes 3.3-4.1× faster. Using the approximate WFA-GPU alignment, the speed increase is even greater, at 15.1-63×, while maintaining 100% accuracy.

When executing WFA-GPU without calculating the alignment path (only calculating the alignment distance), the process is 1.8-4.7× faster than the baseline WFA-GPU. This results in a speed increase of up to 17× compared to the CPU WFA implementation. In addition, we implement an approximated distance-only kernel, which gives a speedup of up to 135.2× compared to the multithreaded CPU WFA implementation.

### 3.3 Evaluation on Real Data

Table 1 compares the performance of WFA-GPU to other state-of-the-art libraries and tools when aligning real datasets (listed in Table S1). Figure 3 illustrates the results of the most relevant datasets and implementations.

**Table 1.**
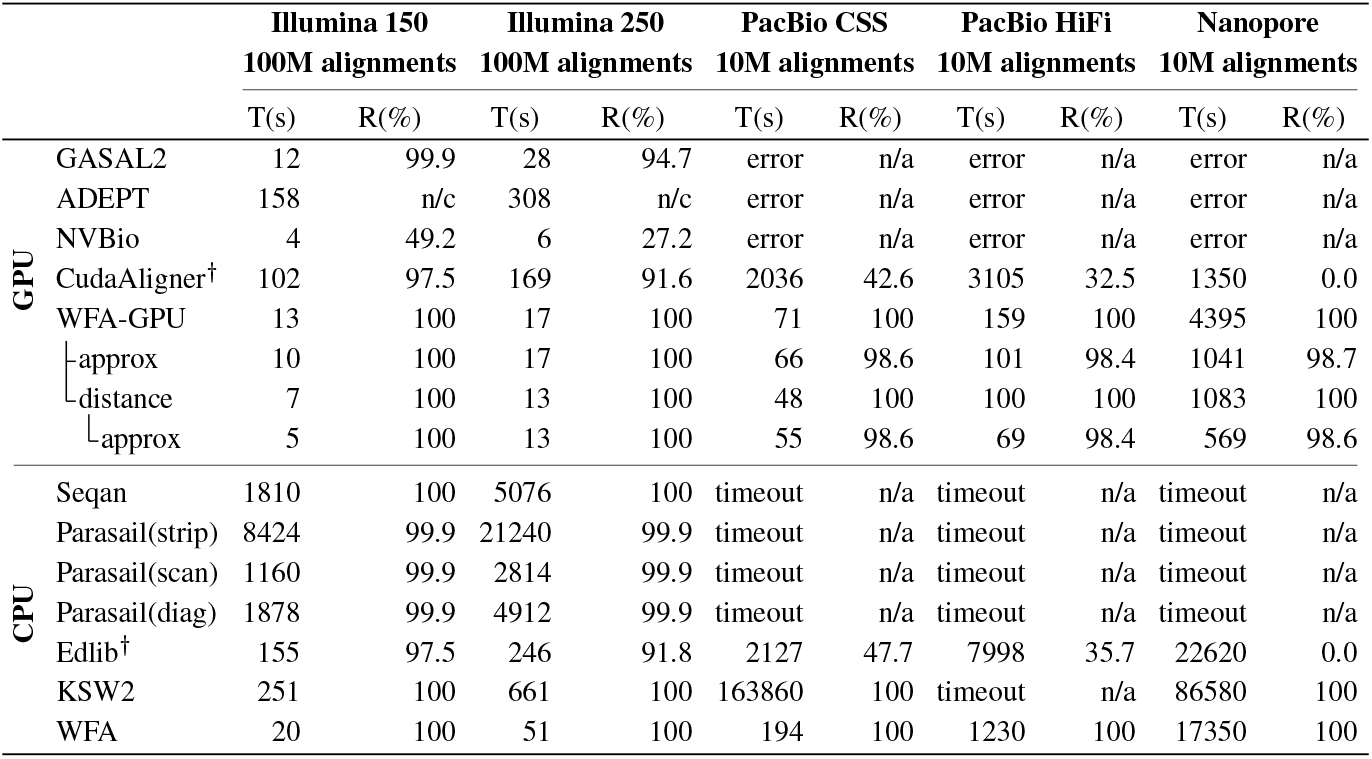
Time (T, in seconds) and recall (R, as a percentage of exact alignments) for real datasets. All CPU executions use 10 threads. ^†^Implementations that can only produce edit-distance alignments. Executions taking more than 48 hours are marked as timeout.

**Fig. 3:**
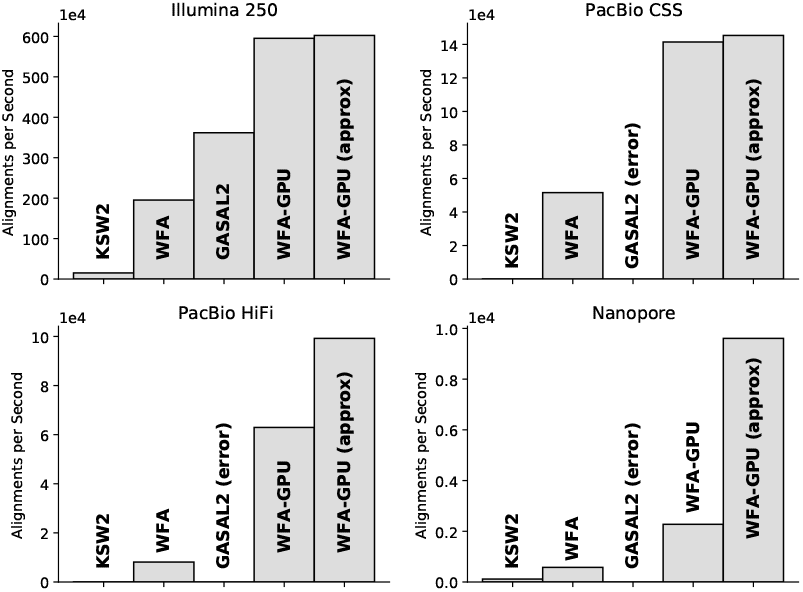
Alignments per second obtained with the most performant CPU and GPU gap-affine implementations compared with WFA-GPU.

For the case of aligning high-quality short sequences, like those produced by Illumina sequencers, NVBio delivers the fastest results at the expense of scoring low in recall (only 49.2% and 27.2% of the alignments are correct). GASAL2 delivers similar performance to WFA-GPU when aligning the Illumina 150 dataset and is 1.7× slower when aligning Illumina 250 (which has slightly longer sequences). Compared to the original WFA, which achieves the best execution time among all CPU libraries, WFA-GPU is 1.5-3× faster. Compared to other CPU libraries, WFA-GPU obtains remarkable speedups (up to 1264× with respect to Parasail). On these datasets (that have a small nominal error), using WFA-GPU approximate alignment has little effect, providing an additional speed increase of only 1.3× (with a total speedup of 2-3×). Computing only the alignment distance (distance-only kernel) is 1.4-2× faster than computing the whole alignment, giving an overall speedup of 3.2-4.2× compared with the multithreaded CPU WFA implementation.

Using PacBio sequences, WFA-GPU achieves a speedup of 2.7× (on PacBio CSS) and 7.7× (on PacBio HiFi) compared to the multithreaded CPU version of the WFA. The speedup raises up to 12.1× if we don’t compute the alignment path (distance-only version). Our implementation outperforms by up to four orders of magnitude other CPU tools and libraries. The only GPU implementation able to finish is CudaAligner, even though it obtains a significantly low recall (less than 50%) while being between 19.5-26.7× slower than our solution. When using WFA-GPU approximate alignment, we obtain an overall speedup of 3.5-17.9× for the CSS and HiFi datasets, respectively. Combining the approximated approach and the distance-only kernel, the speedup goes up to 17.9×.

Regarding the Nanopore dataset, which consists of large sequences with a high error rate, WFA-GPU is 4× faster than the CPU implementation of WFA and 16× faster when using the distance-only version. In comparison, CudaAligner is 3× faster for this execution, but it generates incorrect results (0% recall). When compared to other CPU libraries, WFA-GPU is up to 19.7× faster. However, this speedup is not as high as on previous datasets because alignments with a high nominal error represent the worst-case scenario for the WFA algorithm. Despite this, a high error rate presents a good opportunity for approximated methods to improve execution times. In particular, the WFA-GPU approximated approach obtains a speed increase of 16× compared with the multithreaded WFA CPU version. Only computing the alignment distance while using approximate WFA-GPU results in a speedup of 30.2×.

In general, other GPU implementations either fail to scale with increasing sequence lengths, drop enormously in the recall, or simply fail to execute. Although CudaAligner delivers good performance results, it is bounded to produce edit-distance alignments that fail to capture the biological insights that gap-affine models do. While traditional DP-based algorithms fail to scale with sequence length, WFA-based implementations demonstrate to scale with increasing error rates and lengths. In practice, DP-based implementations require impractical execution times. As opposed, our WFA-GPU implementation delivers good performance results, even when aligning long and noisy sequences. Additionally, our approximate alignment method for WFA can provide additional speedups with little accuracy compromise (less than 2% of sub-optimal alignments reported).

## 4 Discussion

Future advances in sequencing technologies and genomics present critical challenges to current bioinformatics methods and tools. This situation calls for improved methods and tools that can scale with increasing production yields and sequence lengths. Current HPC computing relies on GPUs as successful hardware accelerators for computing-intensive applications in many areas of research. This work presents the first GPU-based tool for sequence alignment based on the efficient WFA algorithm. We proposed algorithmic adaptations and optimizations of the WFA to effectively parallelize the alignment task, exploiting the high-performance capabilities of modern GPU cards.

We demonstrate the benefits of WFA-GPU compared to other state-of-the-art CPU and GPU tools and libraries. Our WFA-GPU implementation performs up to 29× faster than other GPU tools, and up to four orders of magnitude faster than DP-based CPU libraries. Compared to the WFA CPU implementation (fastest CPU library to date) we obtain speedups between 1.5-7.7× on real datasets, without any accuracy loss. When computing only the distance, we get an extra speedup up to 4×. Using our WFA-GPU approximated alignment strategy to align long and noisy sequences, our method reaches a maximum speedup of 16.7× compared to the original WFA CPU, retaining a 98.6% accuracy. To the best of our knowledge, WFA-GPU is the only GPU-based pairwise aligner capable of producing exact gap-affine alignments for long-sequencing datasets, like PacBio HiFi or Oxford Nanopore, in reasonable time using commodity GPU devices.

With the advent of improved sequencing technologies and more sophisticated genomic studies, WFA-GPU offers an accurate, fast, and scalable sequence alignment solution that effectively exploits the massive computing capabilities of modern GPU devices. Therefore, we hope that WFA-GPU will become a valuable and practical addition to the bioinformatics toolkit that supports efficient research in future genome analysis.

## References

(2022). Cudaaligner. https://github.com/clara-parabricks/ GenomeWorks. Accessed: 2022-04-06.

(2022). Wfa2 library. https://github.com/smarco/WFA2-lib. xAccessed:d2022-04-09.

Ahmed, N., Lévy, J., Ren, S., Mushtaq, H., Bertels, K., and Al-Ars, Z. (2019). Gasal2: a gpu accelerated sequence alignment library for high-throughput ngs data. BMC Bioinformatics, 20(1), 520.

Alser, M., Hassan, H., Xin, H., Ergin, O., Mutlu, O., and Alkan, C. (2017). Gatekeeper: a new hardware architecture for accelerating pre-alignment in dna short read mapping. Bioinformatics, 33(21), 3355–3363.

Alser, M., Hassan, H., Kumar, A., Mutlu, O., and Alkan, C. (2019). Shouji: a fast and efficient pre-alignment filter for sequence alignment. Bioinformatics, 35(21), 4255–4263.

Awan, M. G., Deslippe, J., Buluc, A., Selvitopi, O., Hofmeyr, S., Oliker, L., and Yelick, K. (2020). Adept: a domain independent sequence alignment strategy for gpu architectures. BMC bioinformatics, 21(1), 1–29.

Baeza-Yates, R. (1989).Efficient text searching. University of Waterloo.

Baeza-Yates, R. A. (1992). Text-retrieval: Theory and practice. In IFIP Congress (1), volume 12, pages 465–476. Citeseer.

Chacón, A., Marco-Sola, S., Espinosa, A., Ribeca, P., and Moure, J. C. (2014). Thread-cooperative, bit-parallel computation of levenshtein distance on gpu. In Proceedings of the 28th ACM international conference on Supercomputing, pages 103–112.

Daily, J. (2016). Parasail: Simd c library for global, semi-global, and local pairwise sequence alignments. BMC bioinformatics, 17(1), 1–11.

Döring, A., Weese, D., Rausch, T., and Reinert, K. (2008). Seqan an efficient, generic c++ library for sequence analysis. BMC bioinformatics, 9(1), 1–9.

Durbin, R., Eddy, S. R., Krogh, A., and Mitchison, G. (1998). Biological sequence analysis: probabilistic models of proteins and nucleic acids. Cambridge University Press.

Eizenga, J. M. and Paten, B. (2022). Improving the time and space complexity of the wfa algorithm and generalizing its scoring. bioRxiv, pages 2022–01.

Farrar, M. (2007). Striped smith–waterman speeds database searches six times over other simd implementations. Bioinformatics, 23(2), 156–161.

Hwu, W.-M. W. (2011). GPU computing gems emerald edition. Morgan Kaufmann Publishers Inc.

Jones, N. C., Pevzner, P. A., and Pevzner, P. (2004). An introduction to bioinformatics algorithms. MIT press.

Koren, S., Walenz, B. P., Berlin, K., Miller, J. R., Bergman, N. H., and Phillippy,M. (2017). Canu: scalable and accurate long-read assembly via adaptive k-mer weighting and repeat separation. Genome Research, 27(5), 722–736.

Li, H. (2013). Aligning sequence reads, clone sequences and assembly contigs with bwa-mem. arXiv preprint 1303.3997.

Li, H. (2018). Minimap2: pairwise alignment for nucleotide sequences. Bioinformatics, 34(18), 3094–3100.

Lin, C.-H., Li, J.-C., Liu, C.-H., and Chang, S.-C. (2017). Perfect hashing based parallel algorithms for multiple string matching on graphic processing units. IEEE Transactions on Parallel and Distributed Systems, 28(9), 2639–2650.

Lindegger, J., Cali, D. S., Alser, M., Gómez-Luna, J., and Mutlu, O. (2022). Algorithmic improvement and gpu acceleration of the genasm algorithm. arXiv preprint arXiv:2203.15561.

Marco-Sola, S., Sammeth, M., Guigó, R., and Ribeca, P. (2012). The gem mapper: fast, accurate and versatile alignment by filtration. Nature Methods, 9(12), 1185–1188.

Marco-Sola, S., Moure, J. C., Moreto, M., and Espinosa, A. (2021). Fast gap-affine pairwise alignment using the wavefront algorithm. Bioinformatics, 37(4), 456–463.

McKenna, A., Hanna, M., Banks, E., Sivachenko, A., Cibulskis, K., Kernytsky, A., Garimella, K., Altshuler, D., Gabriel, S., Daly, M., et al. (2010). The genome analysis toolkit: a mapreduce framework for analyzing next-generation dna sequencing data. Genome Research, 20(9), 1297–1303.

Myers, E. W. (1986). An o(nd) difference algorithm and its variations. Algorithmica, 1(1-4), 251–266.

Myers, G. (1999). A fast bit-vector algorithm for approximate string matching based on dynamic programming. Journal of the ACM, 46(3), 395–415.

Navarro, G. (1997). A partial deterministic automaton for approximate string matching. Department of Computer Science, University of Chile.

Owens, J. D., Houston, M., Luebke, D., Green, S., Stone, J. E., and Phillips, J. C. (2008). Gpu computing. Proceedings of the IEEE, 96(5), 879–899.

Pantaleoni J, S. N. (2015). Nvbio. https://nvlabs.github.io/nvbio. xAccessed: 2021-09-15.

Petersen, L. M., Martin, I. W., Moschetti, W. E., Kershaw, C. M., and Tsongalis, G. J. (2019). Third-generation sequencing in the clinical laboratory: exploring the advantages and challenges of nanopore sequencing. Journal of Clinical Microbiology, 58(1), e01315–19.

Rodríguez-Martín, B., Palumbo, E., Marco-Sola, S., Griebel, T., Ribeca, P., Alonso, G., Rastrojo, A., Aguado, B., Guigó, R., and Djebali, S. (2017). Chimpipe: accurate detection of fusion genes and transcription-induced chimeras from rna-seq data. BMC genomics, 18(1), 1–17.

Rognes, T. and Seeberg, E. (2000). Six-fold speed-up of smith–waterman sequence database searches using parallel processing on common microprocessors. Bioinformatics, 16(8), 699–706.

Sellers, P. H. (1980). The theory and computation of evolutionary distances: pattern recognition. Journal of Algorithms, 1(4), 359–373.

Simpson, J. T., Wong, K., Jackman, S. D., Schein, J. E., Jones, S. J., and Birol, I. (2009). Abyss: a parallel assembler for short read sequence data. Genome Research, 19(6), 1117–1123.

Šošić, M. and Šikić, M. (2017). Edlib: a c/c++ library for fast, exact sequence alignment using edit distance. Bioinformatics, 33(9), 1394–1395.

Suzuki, H. and Kasahara, M. (2017). Acceleration of nucleotide semi-global alignment with adaptive banded dynamic programming. BioRxiv, page 130633.

Suzuki, H. and Kasahara, M. (2018). Introducing difference recurrence relations for faster semi-global alignment of long sequences. BMC bioinformatics, 19(1), 33–47.

Ukkonen, E. (1985). Finding approximate patterns in strings. Journal of Algorithms, 6(1), 132–137.

Wozniak, A. (1997). Using video-oriented instructions to speed up sequence comparison. Bioinformatics, 13(2), 145–150.

Wu, S. and Manber, U. (1992). Fast text searching: allowing errors. Communications of the ACM, 35(10), 83–91.

Zeni, A., Guidi, G., Ellis, M., Ding, N., Santambrogio, M. D., Hofmeyr, S., Buluç, A., Oliker, L., and Yelick, K. (2020). Logan: High-performance gpu-based x-drop long-read alignment. In 2020 IEEE International Parallel and Distributed Processing Symposium (IPDPS),pages 462–471. IEEE.

Zhao, M., Lee, W.-P., Garrison, E. P., and Marth, G. T. (2013). Ssw library: an simd smith-waterman c/c++ library for use in genomic applications. PloS one, 8(12).

Zorita, E., Cusco, P., and Filion, G. J. (2015). Starcode: sequence clustering based on all-pairs search. Bioinformatics, 31(12), 1913–1919.

Zou, Q., Lin, G., Jiang, X., Liu, X., and Zeng, X. (2020). Sequence clustering in bioinformatics: an empirical study. Briefings in bioinformatics, 21(1), 1–10.

