## Supplementary material for "WFA-GPU: Gap-affine pairwise alignment using GPUs": Tables S1 and S2

### 1 Experimental Datasets

| Dataset | Sample | No. pairs | Seq. Length (bps) min | avg | max |
|---|---|---|---|---|---|
| Illumina 150 | HG002† | 100M | 134 | 148 | 162 |
| Illumina 250 | HG002† | 100M | 140 | 248 | 275 |
| PacBio CSS | HG002† | 10M | 263 | 9561 | 18317 |
| PacBio HiFi | HG002§ | 10M | 201 | 12847 | 24640 |
| Nanopore | ERR3279200<br>ERR3279201 | 10M | 104 | 6249 | 21708 |

Table S1: Description of the real datasets used in the experimental evaluation. †Datasets obtained from NIST's Genome in a Bottle (GIAB) project can be found at `https://github.com/genome-in-a-bottle/giab_data_indexes`. §Datasets obtained from PrecisionFDA Truth Challenge V2 can be found at `https://precision.fda.gov/challenges/10`.

### 2 Experimental Evaluation on Simulated Data

| | | **100 Million alignments × 150bps** | | | | | | **10 Million alignments × 1Kbps** | | | | | | **100K alignments × 10Kbps** | | | | | |
|---|---|---|---|---|---|---|---|---|---|---|---|---|---|---|---|---|---|---|---|
| | | **e=2%** | | **e=5%** | | **e=10%** | | **e=2%** | | **e=5%** | | **e=10%** | | **e=2%** | | **e=5%** | | **e=10%** | |
| | | T(s) | R(%) | T(s) | R(%) | T(s) | R(%) | T(s) | R(%) | T(s) | R(%) | T(s) | R(%) | T(s) | R(%) | T(s) | R(%) | T(s) | R(%) |
| **GPU** | GASAL2 | 13 | 98.3 | 14 | 95.8 | 14 | 93.0 | 44 | 84.6 | 44 | 64.7 | 45 | 52.4 | 3219 | 47.0 | 3224 | 36.8 | 3229 | 34.1 |
| | ADEPT | 161 | n/c | 161 | n/c | 161 | n/c | error | n/a | error | n/a | error | n/a | error | n/a | error | n/a | error | n/a |
| | NVBio | 6 | 58.2 | 6 | 6.1 | 6 | 0.0 | error | n/a | error | n/a | error | n/a | error | n/a | error | n/a | error | n/a |
| | CudaAligner† | 113 | 95.3 | 113 | 66.1 | 117 | 25.7 | 86 | 9.1 | 86 | 1.6 | 86 | 0.3 | 20 | 0.9 | 20 | 0.0 | 20 | 0.0 |
| | WFA-GPU | 11 | 100 | 18 | 100 | 27 | 100 | 6 | 100 | 20 | 100 | 78 | 100 | 4 | 100 | 21 | 100 | 71 | 100 |
| | ├approx | 11 | 100 | 18 | 100 | 27 | 100 | 6 | 100 | 11 | 100 | 21 | 100 | 1 | 100 | 2 | 100 | 4 | 100 |
| | └distance | 6 | 100 | 8 | 100 | 13 | 100 | 3 | 100 | 9 | 100 | 32 | 100 | 2 | 100 | 6 | 100 | 16 | 100 |
| | └approx | 6 | 100 | 7 | 100 | 10 | 100 | 3 | 100 | 4 | 100 | 7 | 100 | 1 | 100 | 1 | 100 | 2 | 100 |
| **CPU** | Seqan | 2028 | 100 | 2028 | 100 | 2190 | 100 | 9066 | 100 | 9288 | 100 | 9774 | 100 | 9912 | 100 | 10098 | 100 | 10440 | 100 |
| | Parasail(strip) | 8436 | 100 | 8358 | 100 | 8238 | 100 | 32274 | 100 | 32178 | 100 | 31842 | 100 | 33408 | 100 | 33306 | 100 | 32760 | 100 |
| | Parasail(scan) | 1197 | 100 | 1183 | 100 | 1197 | 100 | 3882 | 100 | 3902 | 100 | 3911 | 100 | 4869 | 100 | 4857 | 100 | 4825 | 100 |
| | Parasail(diag) | 1908 | 100 | 1914 | 100 | 1916 | 100 | 7482 | 100 | 7488 | 100 | 7488 | 100 | 7686 | 100 | 7692 | 100 | 7698 | 100 |
| | Edlib† | 210 | 95.9 | 202 | 69.6 | 200 | 30.0 | 118 | 67.6 | 118 | 9.5 | 135 | 0.0 | 36 | 1.8 | 38 | 0.0 | 43 | 0.0 |
| | KSW2 | 298 | 100 | 306 | 100 | 317 | 100 | 876 | 100 | 882 | 100 | 882 | 100 | 2028 | 100 | 2034 | 100 | 2040 | 100 |
| | WFA | 20 | 100 | 52 | 100 | 77 | 100 | 34 | 100 | 70 | 100 | 199 | 100 | 15 | 100 | 86 | 100 | 277 | 100 |

Table S2: Time (T, in seconds) and recall (R, as a percentage of exact alignments) for simulated datasets with different error rates (e). All CPU executions use 10 threads. †Implementations can only produce edit-distance alignments.
